# Prophages are Infrequently Associated With Antibiotic Resistance in *Pseudomonas aeruginosa* Clinical Isolates

**DOI:** 10.1101/2024.06.02.595912

**Authors:** Tony H. Chang, Julie D. Pourtois, Naomi L. Haddock, Daisuke Furkuawa, Kate E. Kelly, Derek F. Amanatullah, Elizabeth Burgener, Carlos Milla, Niaz Banaei, Paul L. Bollyky

## Abstract

Lysogenic bacteriophages can integrate their genome into the bacterial chromosome in the form of a prophage and can promote genetic transfer between bacterial strains *in vitro*. However, the contribution of lysogenic phages to the incidence of antimicrobial resistance (AMR) in clinical settings is poorly understood. Here, in a set of 186 clinical isolates of *Pseudomonas aeruginosa* collected from respiratory cultures from 82 patients with cystic fibrosis (CF), we evaluate the links between prophage counts and both genomic and phenotypic resistance to six anti-pseudomonal antibiotics: tobramycin, colistin, ciprofloxacin, meropenem, aztreonam, and piperacillin-tazobactam. We identified 239 different prophages in total. We find that *P. aeruginosa* isolates contain on average 3.06 +/- 1.84 (SD) predicted prophages. We find no significant association between the number of prophages per isolate and the minimum inhibitory concentration (MIC) for any of these antibiotics. We then investigate the relationship between particular prophages and AMR. We identify a single lysogenic phage associated with phenotypic resistance to the antibiotic tobramycin and, consistent with this association, we observe that AMR genes associated with resistance to tobramycin are more likely to be found when this prophage is present. However we find that they are not encoded directly on prophage sequences. Additionally, we identify a single prophage statistically associated with ciprofloxacin resistance but do not identify any genes associated with ciprofloxacin phenotypic resistance. These findings suggest that prophages are only infrequently associated with the AMR genes in clinical isolates of *P. aeruginosa*.

**Importance:** Antibiotic-resistant infections of *Pseudomonas aeruginosa*, a leading pathogen in patients with Cystic Fibrosis (CF) are a global health threat. While lysogenic bacteriophages are known to facilitate horizontal gene transfer, their role in promoting antibiotic resistance in clinical settings remains poorly understood. In our analysis of 186 clinical isolates of *P. aeruginosa* from CF patients, we find that prophage abundance does not predict phenotypic resistance to key antibiotics but that specific prophages are infrequently associated with tobramycin resistance genes. In addition, we do not find antimicrobial resistance (AMR) genes encoded directly on prophages. These results highlight that while phages can be associated with AMR, phage-mediated AMR transfer may be rare in clinical isolates and difficult to identify. This work is important for future efforts on mitigating AMR in Cystic Fibrosis and other vulnerable populations affected by *Pseudomonas aeruginosa* infections and advances our understanding of bacterial-phage dynamics in clinical infections.

## Observation

Mobile genetic elements such as plasmids, transposable elements and integrons play an important role in the dissemination of antimicrobial resistance (AMR) genes across bacterial populations via horizontal gene transfer. Plasmids, in particular, often carry multiple resistance genes that confer multidrug resistance to their bacterial hosts (Carattoli, 2013; Partridge et al., 2018). Transposable elements, including insertion sequences and transposons, likewise facilitate the movement of resistance genes and contribute to the spread of AMR (Partridge, 2011; Siguier et al., 2014; Morales et al., 2023).

Bacteriophages (phages), viruses that infect bacteria, represent another form of mobile genetic element occasionally implicated in AMR transfer. Bacteriophages can contribute to the transmission of AMR genes through transduction, a process where bacterial DNA is packaged into phage particles and delivered to new host cells (Colavecchio et al., 2017; Koonin & Makarova, 2013). However, while phage-mediated transduction has been observed in the lab (Zinder & Lederberg 1952, Morse et al 1956, Colavecchio et al., 2017), its relevance to clinical AMR is still unclear.

Prophages are lysogenic phages that remain integrated within the bacterial chromosome until induced to enter the lytic cycle, at which time new phage particles are produced and the cell is lysed (Frost et al., 2005; Touchon et al., 2017). Few phages directly encode AMR genes (Enault et al. 2017, Pfeiffer et al. 2022, Calero-Caceres et al 2019, Colavecchio et al., 2017), however, phages have been shown to contribute to the spread of AMR through either generalized or specialized transduction (Davies & Davies, 2010; Muniesa et al., 2013; Haaber et al., 2016; Enault et al., 2016). Generalized transduction is the mispackaging of bacterial DNA into the phage capsid and is considered relatively uncommon (Volkova et al. 2014). Specialized transduction, on the other hand, occurs when the prophage is excised with some of the adjoining bacterial genetic material (Griffith et al. 2002, Chiang et al. 2019).

Here, we investigate the relationship between the presence of prophages and both genotypic and phenotypic AMR in the setting of clinical isolates of *Pseudomonas aeruginosa* (*Pa*) collected from patients with cystic fibrosis (CF), a genetic disease often associated with chronic bacterial respiratory infections. We use whole-genome sequences and MIC measurements for 186 clinical isolates of *Pa* from 82 patients seen at the Cystic Fibrosis Center at Lucille Packard Children’s Hospital at Stanford University Medical Center. While prophage count is not a direct proxy for transduction, we hypothesize that a high number of prophages may broadly reflect a higher susceptibility to phages, for example through the conservation of common phage receptors, and thus a higher rate of transduction and AMR transmission. Finally, we test directly whether prophages encode for AMR genes.

We first identified all prophages integrated in the genomes of the *Pa* clinical isolates. Prophages were then assigned the same identification number if they were a similar length (within 10% of each other) and showed at least 90% pairwise similarity. We found that the distribution of prophage abundance varied, with most isolates containing between one and four prophages (mean = 3.06+/-1.84) (**Figure S1**). Isolates harboring two prophages were most common, followed closely by those with one prophage. The occurrence of isolates with five or more prophages decreased steeply, with only a few instances of isolates containing up to ten prophages, which was the maximum number we observed. All isolates had at least one prophage.

The majority of prophages appeared only once or a few times, indicating a high diversity of prophages among the isolates, with only a few prophages present in more than 10 isolates (**Figure S1**). The most common prophage was found in 53 isolates out of 186 clinical isolates. This suggests that while a few prophages may be common, the prophage population is predominantly composed of diverse, low-frequency variants. We identified 239 different prophages in total.

We measured the MIC of clinical isolates and categorized them into susceptible, intermediate and resistant isolates for tobramycin, colistin, meropenem, ciprofloxacin, aztreonam, and tazobactam/piperacillin. We found that 38%, 11%, 42%, 56%, 25% and 19% of isolates were either intermediate or resistant to tobramycin, colistin, meropenem, ciprofloxacin, aztreonam, and tazobactam/piperacillin, respectively.

### The number of prophages is not associated with phenotypic resistance to six commonly used antibiotics

We hypothesized that a high number of prophages in an isolate may be used as a proxy for higher levels of transduction and be associated with high phenotypic AMR. We tested this association for six antibiotics commonly-used to treat *Pa* infections in patients with cystic fibrosis: tobramycin, colistin, ciprofloxacin, meropenem, aztreonam, and piperacillin-tazobactam. There was no significant association between the number of prophages per isolate and the minimum inhibitory concentration (MIC) for any of these antibiotics **(Figure 1**, t-test, p > 0.05). This suggests that prophage abundance alone is not a major factor influencing resistance profiles in these bacterial isolates.

**Figure 1:**
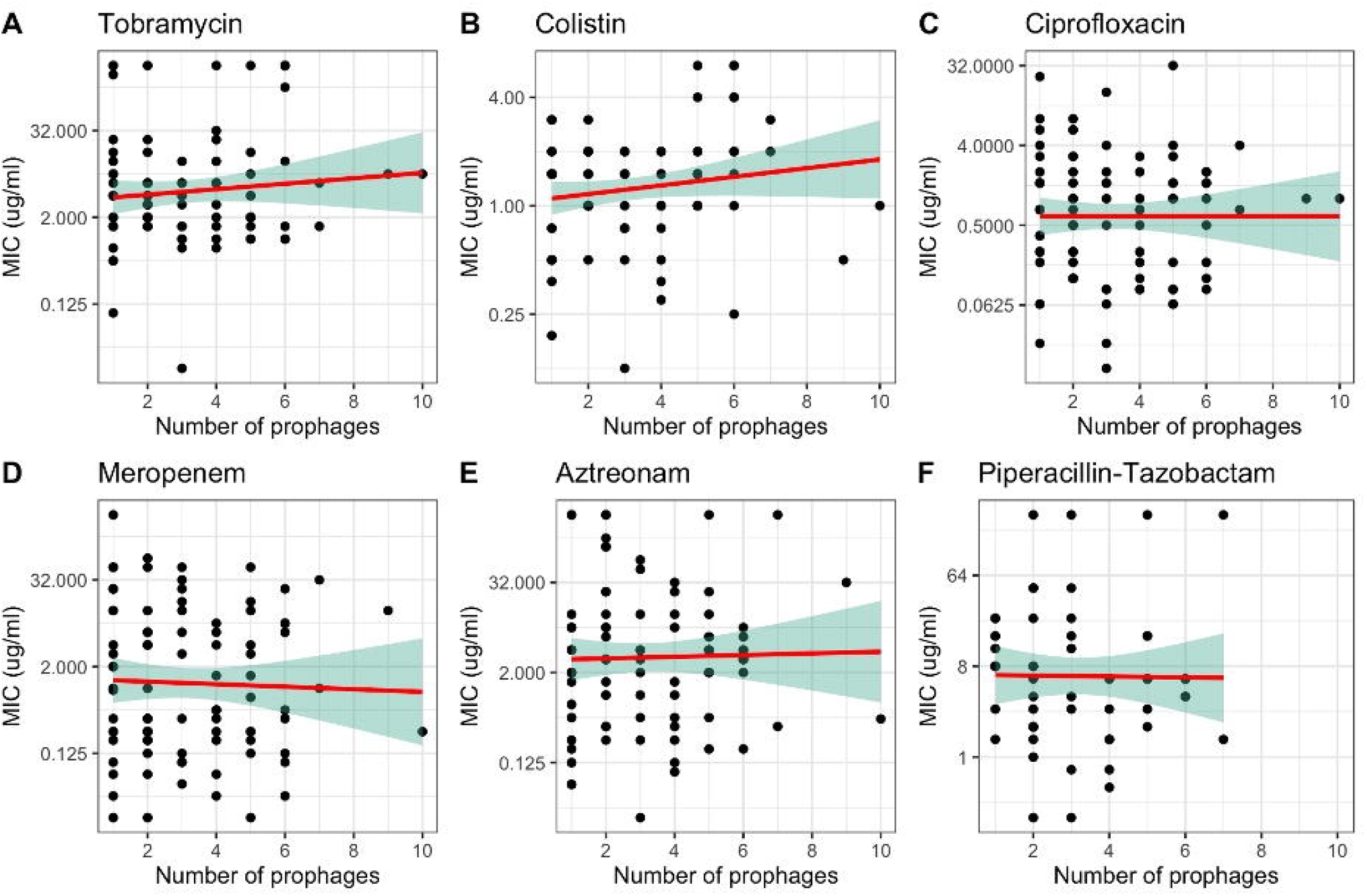
There is no relationship between the Number of Prophages and the Minimum Inhibitory Concentration (MIC) for various antibiotics. **A**. Tobramycin **B**. Colistin **C**. Ciprofloxacin **D**. Meropenem **E**. Aztreonam **F**. Piperacillin-Tazobactam. All y-axis are on a log2 scale. Each plot includes the linear regression line (red) with a shaded 95% confidence interval (green). None of the slope coefficients are significantly different from 0 (p > 0.05).

### Specific prophages are associated with an increase in phenotypic resistance to tobramycin and ciprofloxacin

Most prophages were not present in enough isolates to evaluate their relationship with resistance. We selected the four most prevalent phages (N > 20), which we named vB_Tem_CfSt1-4 and evaluated their association with the MIC for the four antibiotics tobramycin, colistin, ciprofloxacin and meropenem. We found no association between these prophages and resistance to colistin or meropenem (**Figure S2**).

However, we observed a significant increase in tobramycin resistance when prophage vB_Tem_CfSt1 (**Figure 2A**) was present and a significant increase in ciprofloxacin resistance when prophage vB_Tem_CfSt3 was present (**Figure 2B**).

**Figure 2:**
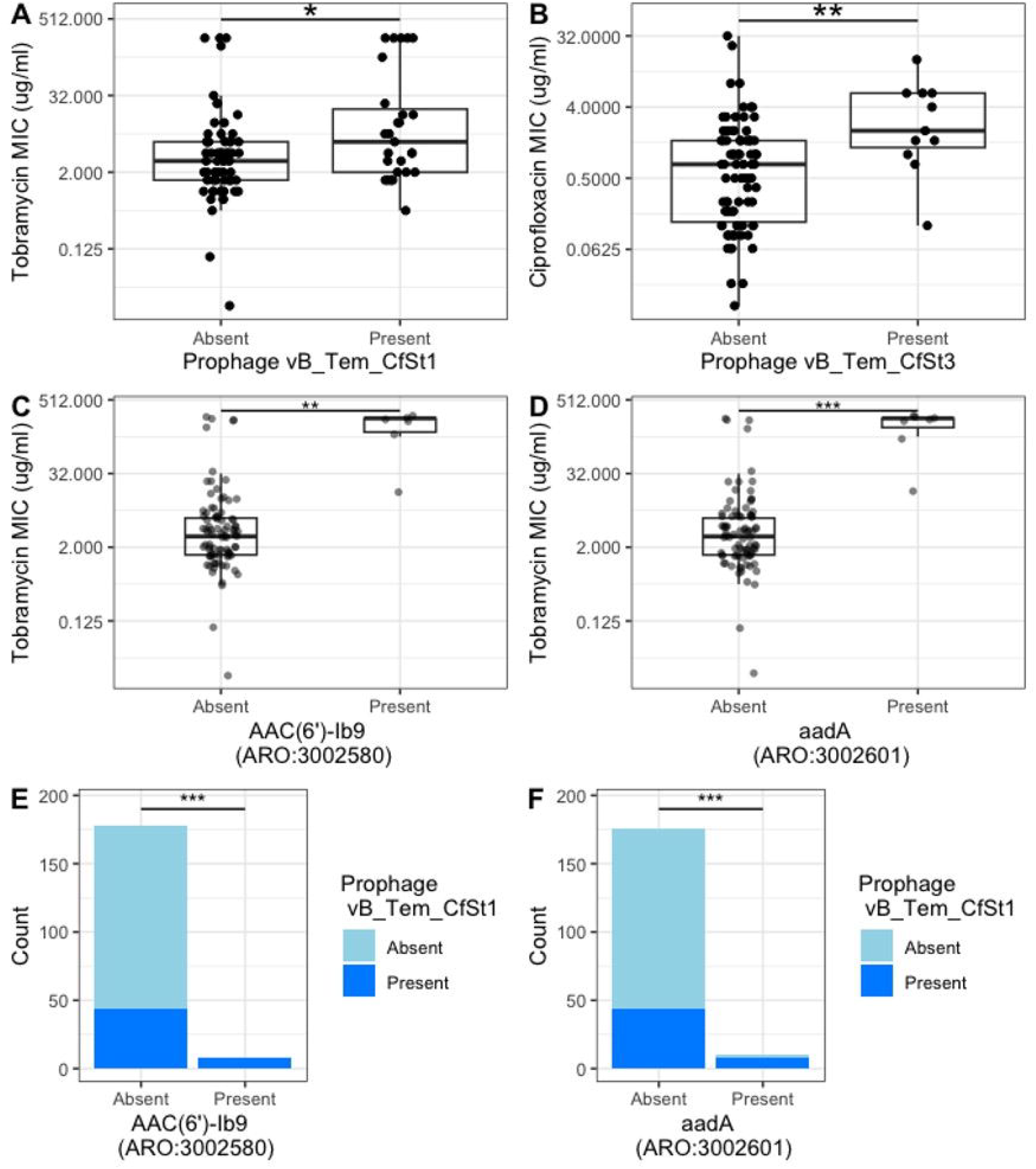
Identification of a pair of prophages that are associated with AMR. We selected the four most prevalent phages which we named vB_Tem_CfSt1-4 and evaluated their association with the MIC of four antibiotics tobramycin, colistin, ciprofloxacin and meropenem. We found no association between these prophages and resistance to colistin or meropenem (Figure S2). However, we observed a significant increase in tobramycin MIC when prophage vB_Tem_CfSt1 (**A**.) was present and a significant increase in ciprofloxacin resistance when prophage vB_Tem_CfSt3 was present (**B**.) ‘*’, ‘** and ‘***’ for p-values smaller than 0.05, 0.01 and 0.001 respectively. **C**. Tobramycin resistance (Tobramycin MIC on a Log2 scale) for isolates with and without resistance genes AAC(6’)-Ib9 (ARO:3002580) **D**. Tobramycin resistance (Log2 Tobramycin MIC) for isolates with and without resistance genes aadA (ARO:3002601) **E**. Presence of temperate phage vB_Tem_CfSt1 for isolates with and without antibiotic resistance genes AAC(6’)-Ib9 **F**. Presence of temperate phage vB_Tem_CfSt1 for isolates with and without antibiotic resistance genes aadA. We use ‘*’, ‘** and ‘***’ for p-values smaller than 0.05, 0.01 and 0.001 respectively.

### Prophage associated with increases in tobramycin MIC is also associated with AMR genes

We next asked if the same prophages were associated with the presence of AMR genes. Among all AMR genes described in the CARD-RGI dataset found in our isolates, only the genes aadA and AAC(6’)-lb9 were significantly associated with higher MIC to tobramycin (Figure 2C & 2D, Wilcoxon test, p < 0.01). Finally, we found that the presence of prophage vB_Tem_CfSt1 was significantly associated with the presence of these genes (**Figure 2E & 2F**, Fisher test, p < 0.01). The genes aadA and AAC(6’)-lb9 both encode aminoglycosides-modifying enzymes (Hollingshead & Vapnek 1985, Mugnier et al, 1998). These enzymes are some of the most widespread mechanisms for tobramycin resistance and are known to be highly mobile (Garneau-Tsodiko & Labby 2016). Other common mechanisms for tobramycin resistance include other mobile genes such as efflux pumps and ribosome methyltransferases. Mutations in the ribosome genes targeted by aminoglycosides are rarer on the other hand.

This is in contrast with ciprofloxacin resistance, which is often acquired through mutations of the A subunit DNA gyrase targeted by quinolones like ciprofloxacin, rather than mobile genetic elements (Hooper et al, 1989). We did not find any genes associated with a higher MIC for ciprofloxacin. Other works also show a lower association between known genotypic markers of resistance and phenotypic resistance for ciprofloxacin compared to other antibiotics like aminoglycosides (Vanstokstraeten et al 2023), which is consistent with our findings. Phage-mediated transduction of AMR genes may thus be more relevant to antibiotics for which mobile genes, rather than point mutations, confer resistance and will be easier to detect for antibiotics with strong genotype to phenotype associations for antibiotic resistance.

### Prophages do not directly encode AMR genes

Finally, we looked for AMR genes in the prophage sequences using the homolog model of the CARD-RGI database. None were identified except for a single occurrence of rsmA, a virulence gene that can also confer antibiotic resistance (Mulcahy et al 2006). This is consistent with previous work showing that AMR genes are rarely encoded directly by phages (Enault et al 2016).

We were not able to determine the relative location of prophages and antibiotic resistance genes due to the small size of many scaffolds containing prophages and AMR genes. The genome reconstructions from the short-read sequencing performed for this work did not allow us to establish the relative locations of prophages and AMR genes. It is thus important to note that the association between prophage vB_Tem_CfSt1 and genes aadA and AAC(6’)-lb9 does not imply causation and could arise from phylogenetic correlation for example. In addition, our findings are limited by low prevalence for many prophages and AMR genes, the majority of which were only found in one isolate. Very few phages were found in more than twenty isolates and many AMR genes associated with tobramycin resistance were either present in all or very few isolates. The low prevalence of most AMR genes also prevented us from assessing additive effects from the presence of multiple genes.

In summary, we identified a single prophage associated with higher MIC for tobramycin and found that known AMR genes conferring resistance to tobramycin were more likely to be found in isolates infected with that phage. Prophage abundance was not associated with antibiotic resistance. These findings suggest that the spread of AMR through prophages is rare overall. Future work should focus on new phenotype prediction tools to increase sample sizes, as well as long read sequencing to explore causal relationships between phage and AMR through the relative locations of mobile elements in bacterial genomes.

### Sample Collection

From June 2020 to June 2023, 186 *Pseudomonas aeruginosa (Pa)* isolates from respiratory cultures of cystic fibrosis patients were collected at Stanford Hospital under IRB approvals #11197. Samples were biobanked with patient consent and de-identified using unique codes.

### DNA Extraction and Sequencing

DNA was extracted from Pa isolates using the DNeasy Blood and Tissue Kit (Qiagen, 69504) and sequenced on an Illumina NovaSeq (100bp paired-end) and an Illumina NextSeq (150bp paired-end). The extraction involved bacterial lysis, DNA purification, and elution, ensuring high-quality DNA suitable for sequencing. Sequencing reads were quality-checked using FASTQualityControl and trimmed with Trimmomatic 0.39 or ‘trim galore’ to remove Nextera adapters from raw reads sequenced on Illumina NextSeq. Quality reports were assembled using MultiQC55. Trimmed reads were then assembled with SPAdes using –isolate and --cov-cutoff auto (Prjibelski *et al*., 2020).

### Prophage identification

We identified prophages using VIBRANT (Kieft, Zhou and Anantharaman, 2020). We then performed a BLAST search (word size = 28, e-value = 0.005) on all predicted prophages against their own sequences. Prophages were grouped under the same name if they were of similar length (less than 10% difference in length) and showed at least 90% pairwise identity.

### Antibiotic susceptibility testing

Clinical isolates were streaked on LB agar plates and incubated at 37°C overnight. The next day, the colonies were picked and suspended in 2 ml of sterile 0.85% saline solution. The inoculum was vortexed and its turbidity was adjusted until it reached 0.5Mc. Farland standard (OD 0.08-0.1). The standardized bacterial suspension was evenly streaked onto 150mm Mueller Hinton agar plates (Thermo Fisher Scientific) using a sterile cotton swab. The plates were allowed to dry for 15 min and the E test strips (Fisher Scientific) were placed onto the plates and incubated at 37°C. The MICs were read after 16-20 hrs. MICs were obtained for 93 clinical isolates.

### Identification of AMR genes

Antibiotic resistance genes were identified with a BLAST search (-word size 28 -culling_limit 1 evalue 0.005) of bacterial genomes against the homolog model of the CARD-RGI database. This model includes sequences that are determinants of resistance without mutation.

### Statistical Analysis

Statistical analyses were performed using R. Tests used include t-tests for linear regression, Wilcoxon tests and fisher tests. Sample sizes were not predetermined. Investigators were blinded during DNA extraction, library preparation, and phage genome annotation, but not during the final analysis.

## Acknowledgements

T.C. discloses support from the Cystic Fibrosis grant student traineeship and Stanford Bio-X foundation grant. P.L.B. discloses support from National Institutes of Health grant R01 HL148184-01, National Institutes of Health grant R01 AI12492093, National Institutes of Health grant R01 DC019965, Cystic Fibrosis Foundation grant, a Stanford SPARK grant, and a grant from the Emerson Collective. The contents are those of the authors and do not necessarily represent the view of the funding agencies. E.B.B. was supported by the Cystic Fibrosis Foundation (BURGEN23G0 and BURGEN24A0-KB) and National Institutes of Health (1K23HL169902-01).

