## Supplemental Data for "Prophages are Infrequently Associated With Antibiotic Resistance in *Pseudomonas aeruginosa* Clinical Isolates"

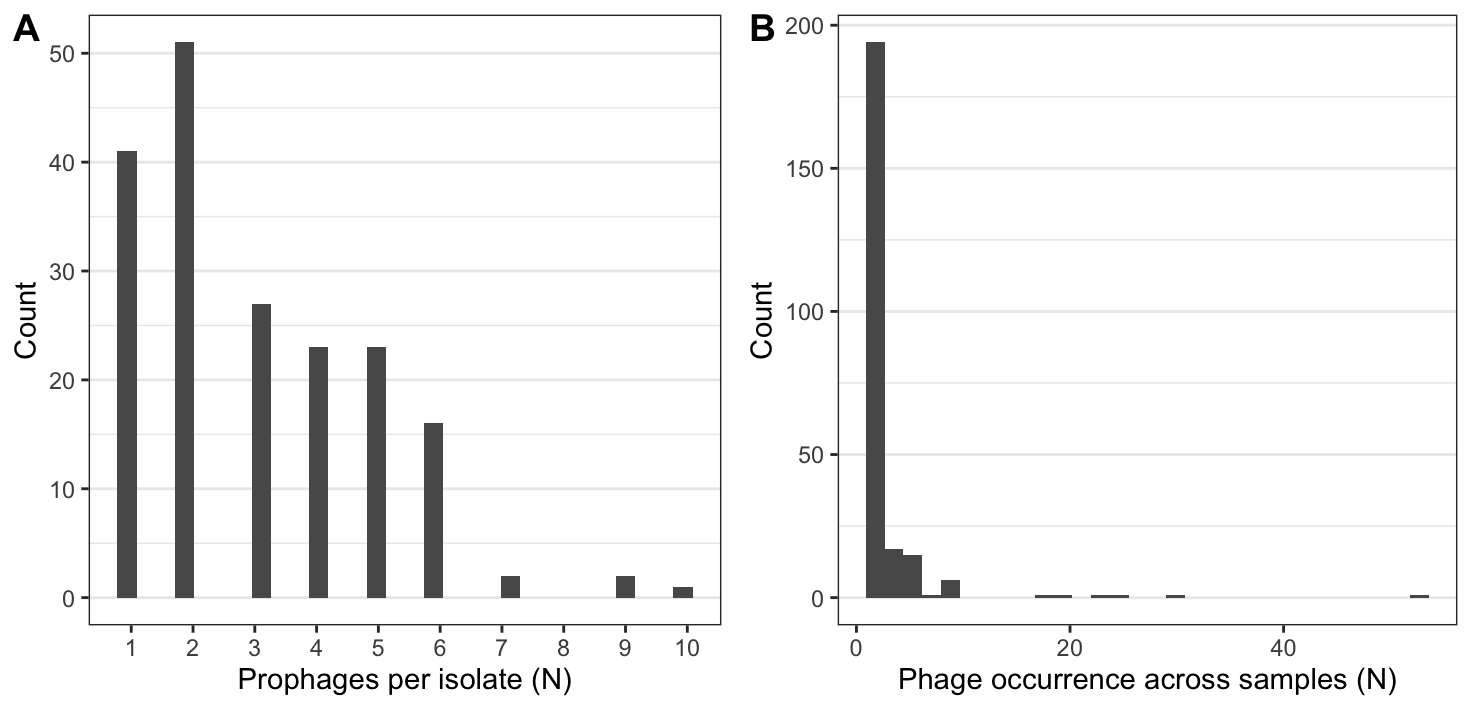

**Figure S1: Prophages are common in clinical samples. A.** Number of prophages per isolate. **B.** Distribution of phage occurrence across the samples. Most phages are present only once or twice with a few phages present many times.

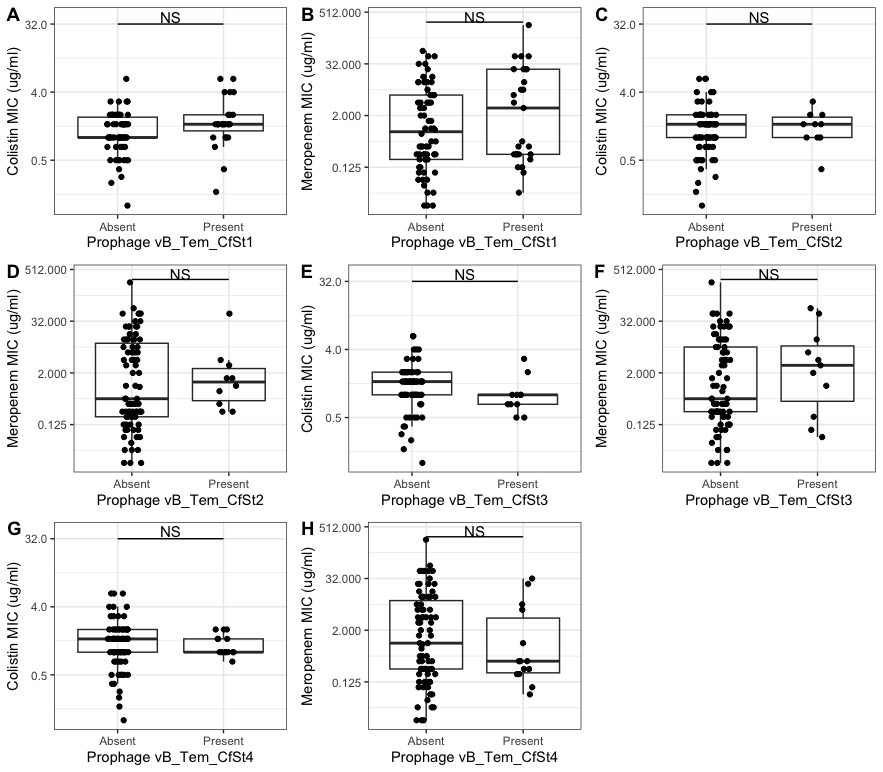

**Figure S2: There was no significant relationship between the presence of any of the 4 most common prophages and phenotypic resistance to colistin (A, C, E, G) or meropenem (B, D, F, H)**.
